# Comprehensive hallmark gene sequence, genomic and structural analysis of *Picornavirales* viruses clarifies new and existing taxa

**DOI:** 10.64898/2026.01.05.697625

**Authors:** Richard Mayne, Donald B. Smith, Katherine Brown, Yan Ping Chen, Andrew E. Firth, Kazuhiko Katayama, Nick J. Knowles, Peter Simmonds

## Abstract

The order *Picornavirales* is a group of highly diverse RNA viruses that includes many pathogens of significance to human and veterinary health, agriculture and the wider environment. However, the wide range of viruses assigned to the order, together with their genomic variability, and the recent description of numerous “picorna-like” viruses derived from metagenomic analyses of environmental samples, challenges the existing taxonomic classification of members of the order and the criteria for their classification. Here, we combine the existing gold standard, hallmark RNA-dependent RNA-polymerase (RdRp) gene sequence-based analysis with helicase sequence-based phylogeny, RdRp structural prediction through the use of ColabFold and Fold Tree, and analysis of coding complete genomes using GRAViTy-V2, to genetically classify 525 *Picornavirales* genomes and recently described “picorna-like” viruses. All analyses were conducted with a bespoke, fully automated pipeline for retrieval of genomes, domain classification and extraction, phylogenetic analysis, and output conditioning, which is available as open-source software. Our results reveal broad support for existing families as well as for six novel families, and 32 new genera. In instances where inconsistencies were found between classification methods, we demonstrate how examination of the pipeline’s output may be used to reconcile differences with respect to the genomic features quantified by the analysis. Automated multimodal taxonomic analysis may save significant resources over manual methods and better define demarcation criteria for families and genera.

## 1 Introduction

As of early 2026, the International Committee on Taxonomy of Viruses (ICTV) has assigned 775 species in eight families in the order *Picornavirales*. However, the community recognises at least 1000 additional unclassified picorna- and picorna-like viruses in publicly available databases, many of which have been assembled from high-throughput metagenomic sequencing studies [36, 27, 5]. Members of the *Picornavirales* are eukaryote-infecting, non-enveloped, positive-strand RNA viruses whose distinguishing genomic characteristic is a three-component ‘replication block’ encoding a helicase (Hel), a proteinase (Pro) and an RNA-dependent RNA-polymerase (RdRp), which are cleaved by an auto-proteolytic mechanism [13]. Additionally, viruses in *Picornavirales* may have a 5’ covalently genome-linked protein (VPg) and a 3’ poly(A) tail, though these features are not present in all constituent families. Possession of an icosahedral virion with characteristic jelly-roll fold capsid proteins and T=3 symmetry was previously another of the order’s defining characteristics, prior to the discovery of exceptions to this rule [11]. Indeed, apart from the commonality in replication-associated proteins, the order is notable for its remarkably high genome organisational diversity, even among other RNA viruses. For example, picorna-and picorna-like viruses may be monopartite or bipartite; mono-, di- or polycistronic; possess a variable number of structural proteins and internal ribosomal entry sites (IRES) [13], and are prone to both recombination events [15] and genome organisation reversal [39]. This variability makes the order challenging to classify taxonomically.

There are currently nine accepted families within *Picornavirales* (Table 1) which include a number of pathogens with significant relevance to human, veterinary, agricultural and environmental health. However, several previously described and characterised viruses are not currently classified to family level, including posaviruses (isolated in porcine faeces, proposed as a new family ‘Posaliviridae’ [8]) and Mayfield virus (pathogen of *Apidae*) [22].

**Table 1:**
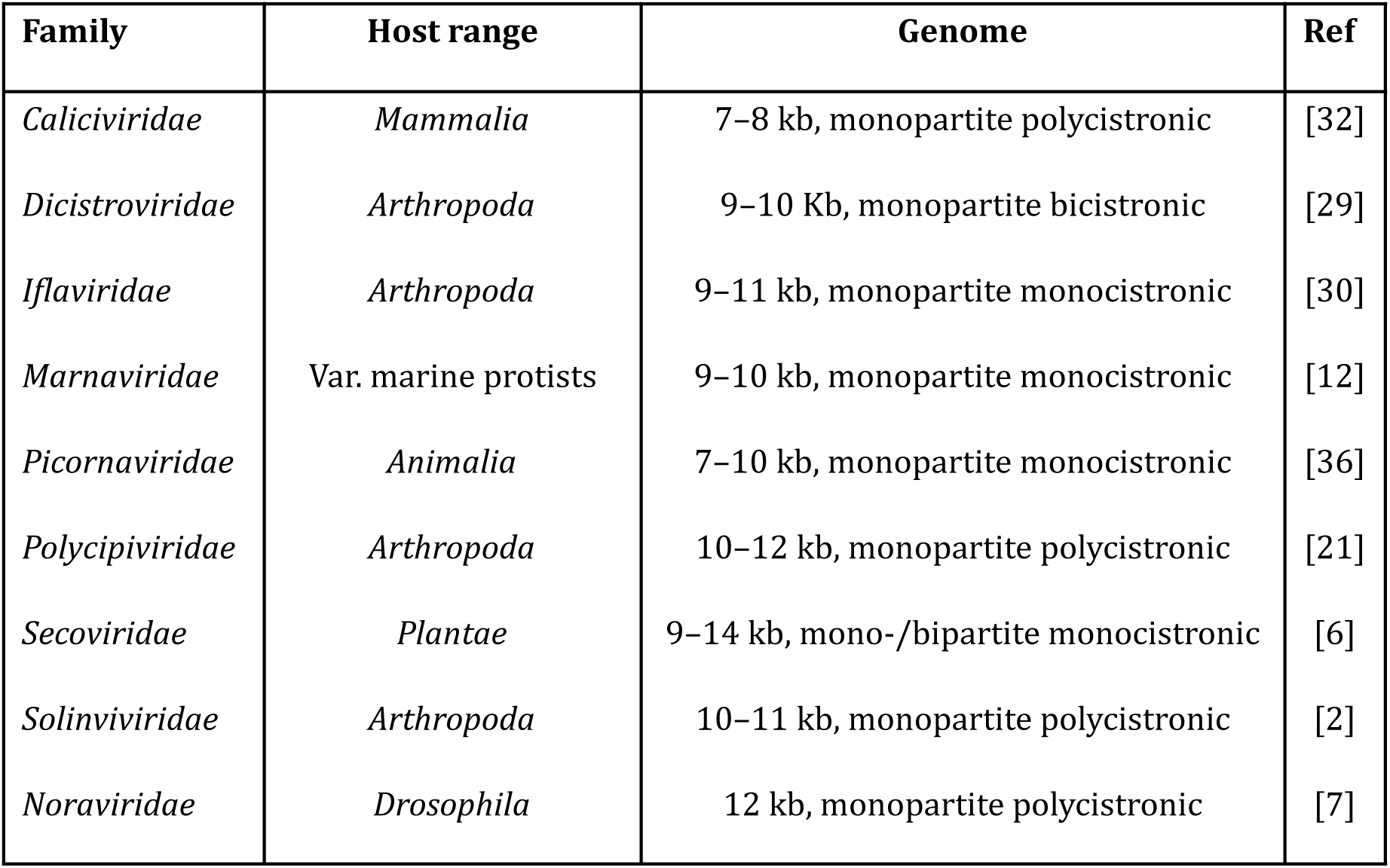
Current family members of the *Picornavirales*.

Taxonomic classification of viruses may only be ratified by the ICTV in cases where the nucleotide sequence for a whole coding region(s) of the genome is known [24, 25]. The gold standard for RNA virus classification at family level and higher ranks remains, however, based on evolutionary relationships of the hallmark RdRp gene, as this genomic component is the only common feature within *Riboviria* (RNA viruses) and therefore uniquely allows virus evolutionary histories to be mapped across the realm. Despite recent international efforts to standardise and adapt RdRp analysis for scale [3], in practice, there is little consensus in the community on what tools and algorithms might be used to achieve this, and manual curation of alignments is commonplace. Such approaches are rational and effective for demonstrating taxonomic relationships between more divergent viruses with common ancestry, but are, however, not scalable and ignore additional genomic, structural and functional (e.g. antigenic) features which are otherwise useful for determining the characteristics and groupings of viruses at the level of genus and species. For example, the widely accepted *Picornaviridae* subfamilies take into consideration not just homologies in the 3CD (Pro-RdRp) domain, but also, structural proteins and genome organisation [38]; meanwhile, *Enterovirus* species are demarcated by a 25% sequence divergence threshold in the major capsid protein VP1 gene.

Generating taxonomies in this manner tends to be extremely time consuming and requires significant amounts of input from subject matter experts. It is likely that, with the power of metagenomic approaches, we will continue to identify more novel viruses within *Picornavirales* and beyond at a rate that greatly exceeds our abilities to classify them with existing methods. This presents an urgent and currently unmet requirement for new, scalable taxonomic methods that can provide more information than the simple linear relationships conveyed by hallmark gene phylogenies.

In this article, we present an analysis of both classified and unclassified picorna- and picorna-like viruses, which combines analysis of two evolutionarily conserved genes (RdRp and Hel), genomic component and organisation similarity, and predicted RdRp protein secondary structure relationships. This multimodal approach builds on our previous work in a recent reclassification and promotion of genera within the *Flaviviridae* to three families within the order *Amarillovirales* [26]. The current study expands this analytical approach through the introduction of an open-source, all-in-one software pipeline which is optimised for use on personal computers and incorporates features such as automated domain extraction and bespoke elements for collating and contrasting discrepancies between methods. We proceed to describe how our results clarify existing and proposed taxonomic groupings for *Picornavirales* and conclude by discussing both limitations and future work in broadening our method to the study of other taxa and at greater scales.

## 2 Methods

The three analytical methodologies were used (Fig. 1) to generate four sets of taxonomic assignments:

a. Alignment and phylogenetic tree generation of both RdRp and Hel domains, using Mafft v7.520 [10] and IQ-TREE 3.0.1 [34].
b. GRAViTy-V2 2.2 [16], which is a framework for discovering and classifying viruses using protein profile hidden Markov models (PPHMMs). A composite Jaccard distance (CJD) metric of shared PPHMM similarity and relative genomic position is used to generate taxonomic assignments from coding-complete genomes.
c. Comparison of structural characteristics of extracted RdRp domains, through ColabFold [18] using a lightweight implementation, ColabFold Local 1.5.5 (https://github.com/YoshitakaMo/localcolabfold) and Fold-Tree (build 02 06 25) [19].

**Figure 1:**
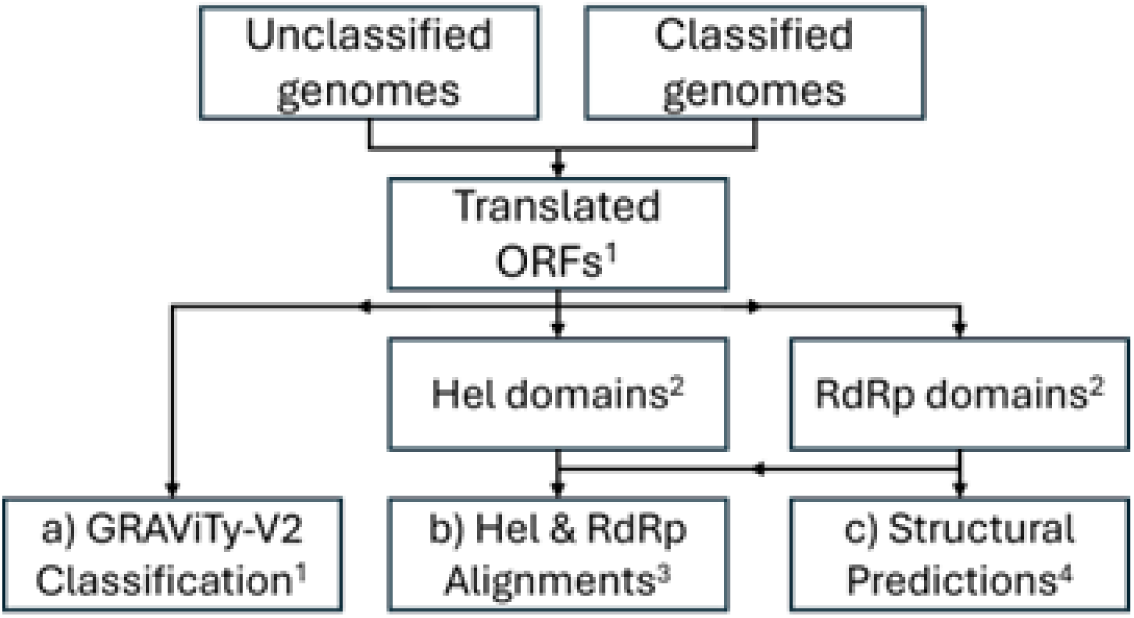
Process flow for three analytical pipelines used (a–c). Tools: ^1^ GRAViTy-V2; ^2^ InterPro Scan; ^3^ Mafft, IQ-Tree3; ^4^ ColabFold Local, Fold Tree.

All analytical methods were executed as part of an automated pipeline, created by the authors, called ‘UniViT’ (https://github.com/Mayne941/UniViT). Output from these tools was in the form of phylogenetic trees rendered with IToL v7 [14], alignments from the hallmark gene methods, and various statistics and PPHMM graphs from GRAViTy-V2. Initial classifications at family and genus levels were generated through a bespoke script as a component of the UniViT pipeline that automatically tabulated leaves from dendrograms for both sequence-based phylogenies and the structural tree and set demarcation boundaries at a fixed depth. These were joined to tabulated GRAViTy-V2 clusters, which were automatically generated by the tool’s classifier module. All classifications were manually checked and disagreements between the phylogenies were recorded.

Trees were compared through the use of Robinson-Foulds metric computed on tree pairs, and through generating tanglegrams. Tanglegrams were rendered in Python 3.10, using a bespoke implementation of the popular algorithm by Nguyen *et al.* [20], over three untangling iterations. Genome diagrams were created using Vectorgraphic_genome v0.1 (https://github.com/donaldsmithictv-cell/Vectorgraphic_genome).

### 2.1 Data acquisition and preprocessing

Previously classified *Picornavirales* sequences were sourced from the ICTV Virus Metadata Resource (VMR, MSL40.v1), which was filtered to one representative example per species (*n* = 244). Unclassified *Picornavirales* sequences were sourced from NCBI Virus, using the filters ‘Unclassified Picornavirales’ and ‘complete genome’ (*n* = 281). As outgroups, additional single representative virus sequences from picorna-like virus families *Potyviridae* (*n* = 13) and *Hypoviridae* (*n* = 2) were included, as were representative members of neighbouring *Amarillovirales* (*n* = 4), all of which were sourced from the VMR. The full dataset is included in SI 1.

Open reading frames (ORFs) were extracted from whole genomes using GRAViTy-V2 and were then input to InterPro Scan [9] to generate protein domain predictions based on the Pfam, PROSITE, PANTHER, SMART and CDD databases. XML outputs from InterPro Scan was parsed with a bespoke script to extract RdRp and Hel amino acid sequences. Sequences were omitted downstream if either or both of the RdRp and Hel domains were not identified.

### 2.2 Hallmark gene alignments

Amino acid multiple sequence alignments were generated for RdRp and Hel sequences output by InterPro Scan, using Mafft with settings:

--localpair --iterate 1000 --leavegappyregion

Alignments were not manually curated but were examined for errors using SSE v1.4 [23]. Maximum-likelihood trees were generated using IQ-TREE using the Q.pfam+F+I+R10 substitution model for both genes and 1000 bootstrap iterations.

### 2.3 GRAViTy-V2

GRAViTy-V2 was run on the dataset described in section 2.1, in single-pass mode using the ‘new classification full’ pipeline. Run parameters and input sequences are included in SI 1. A separate GRAViTy-V2 analysis was performed to include representative single members of each family within *Riboviria*, using two-pass mode but otherwise identical settings to the single-pass run, to reveal relative depths and comparative demarcation boundaries for both orders and families. As an alignment-free taxonomy framework, GRAViTy-V2 represents groups of viruses with no evolutionary relatedness with a CJD distance of 1.0; distances are not empirical measurements of evolutionary distance but do offer a relative scale to compare relatedness between members of homologous groups.

### 2.4 Structural predictions

RdRp sequences were used as input to a structural prediction pipeline, using Colabfold Local. The pipeline then determined the highest confidence models from the output for each structural prediction and predicted Local Distance Difference Test (pLDDT), which were used to generate phylogenetic trees using FoldTree. Structures with amino acid length *>* 20% shorter than median length, or ≤ 80% mean pLDDT confidence scores were omitted from the analysis. A subset of predicted protein structures was manually checked by making family-to-family comparisons (*n* = 25), through using FoldSeek to align and search for homologues in public database. Both pLDDT score for the structure and root mean square deviation (RMSD) of atomic positions for the comparison with the experimentally derived structure were reported. Molecular views were rendered using UCSF ChimeraX [17].

### 2.5 Hardware

All analytical work was conducted on a personal computer running on Ubuntu 22.04 LTS, using an AMD Ryzen 7 7600X 12-core CPU with 32 Gb DDR5 RAM at 6000 MHz. Data were read from and written to an M.2 NVMe SSD at a maximum speed of 3000 MBps. ColabFold was run using an NVidia GeForce RTX 2080 SUPER with 8 Gb GDDR5 RAM and 3072 CUDA cores.

## 3 Results

### 3.1 Comparison of analysis methods

All four analysis methods produced phylogenies that broadly reflected existing family-level classifications, though several rearrangements in genera were observed among methods (Figs. 2-3, SI 2–4). Among the approaches, GRAViTy-V2 classification was most divergent from the RdRP phylogeny (10.5% of classifications disagreed at family level), followed by the Hel phylogeny (7.8%) and the ColabFold structural tree (4.5%) (Table 2). Robinson-Foulds distances between trees ranged from 0.39–0.54, with the Hel phylogeny being the least similar to the RdRP tree. Although genera were generally conserved between RdRp and Hel sequence-based analyses in the larger, more stable families, there were comparatively frequent family-level discrepancies. Although RdRp and Hel domains are anticipated to have co-evolved as a result of their proximity within the replication block [11], the shorter sequence length of the conserved Hel domain confers less resolution for making sequence-based comparisons.

**Figure 2:**
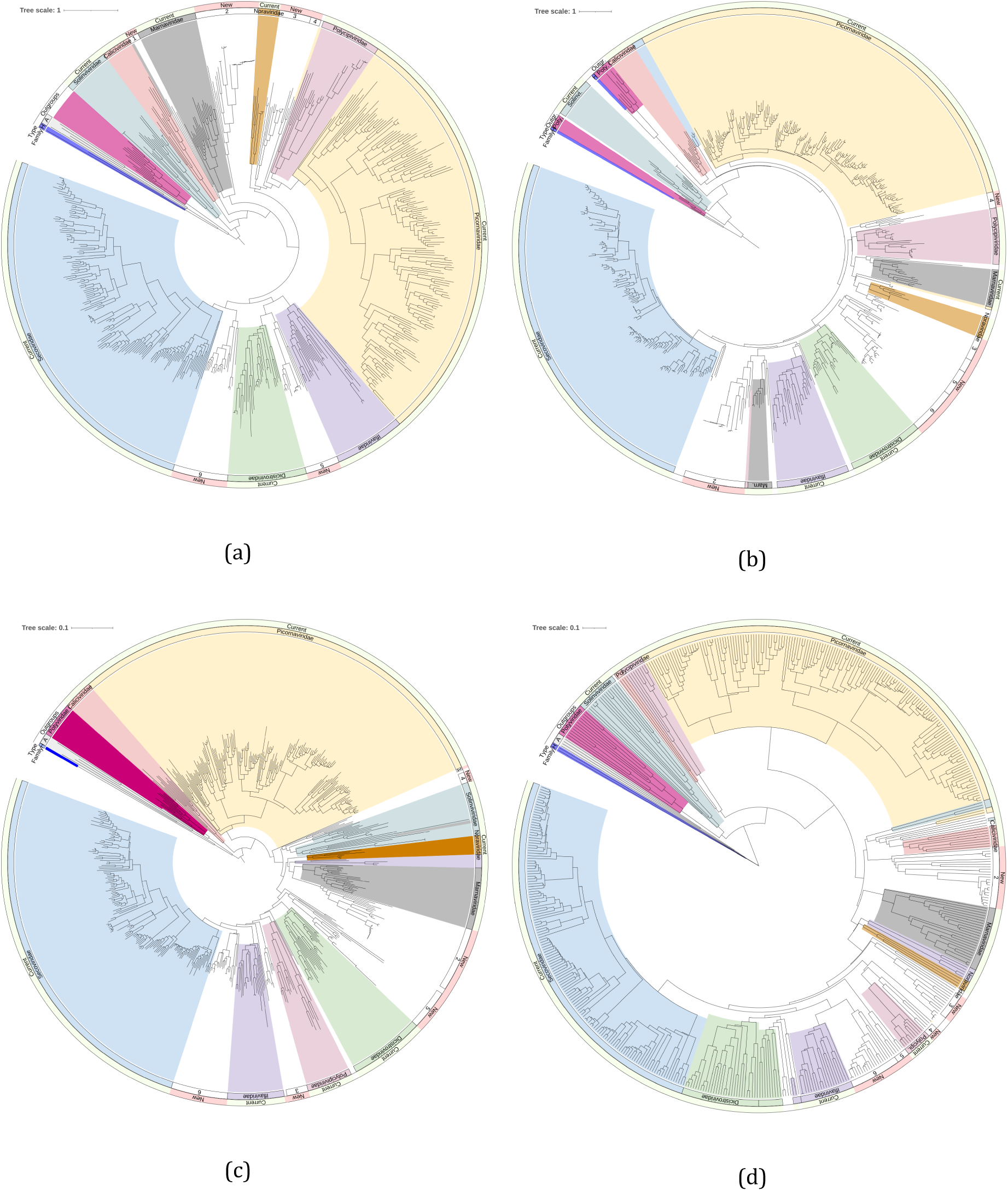
Phylogenetic trees showing conservation of both new and existing family groups between all four analysis methods. Clades are coloured by family where the majority of sequences within a clade correspond to a previously defined family, while numerical annotations indicate proposed new families. (a) RdRp phylogeny. (b) Hel phylogeny. Branches for flaviviruses KC410872, AF160193 trimmed. (c) RdRp structural predictions. Branches trimmed for oat mosaic virus AJ306718 and hypovirus MH766501. (d) GRAViTy-V2 analysis. H.: *Hypoviridae*, A.: *Amarillovirales*.

**Figure 3:**
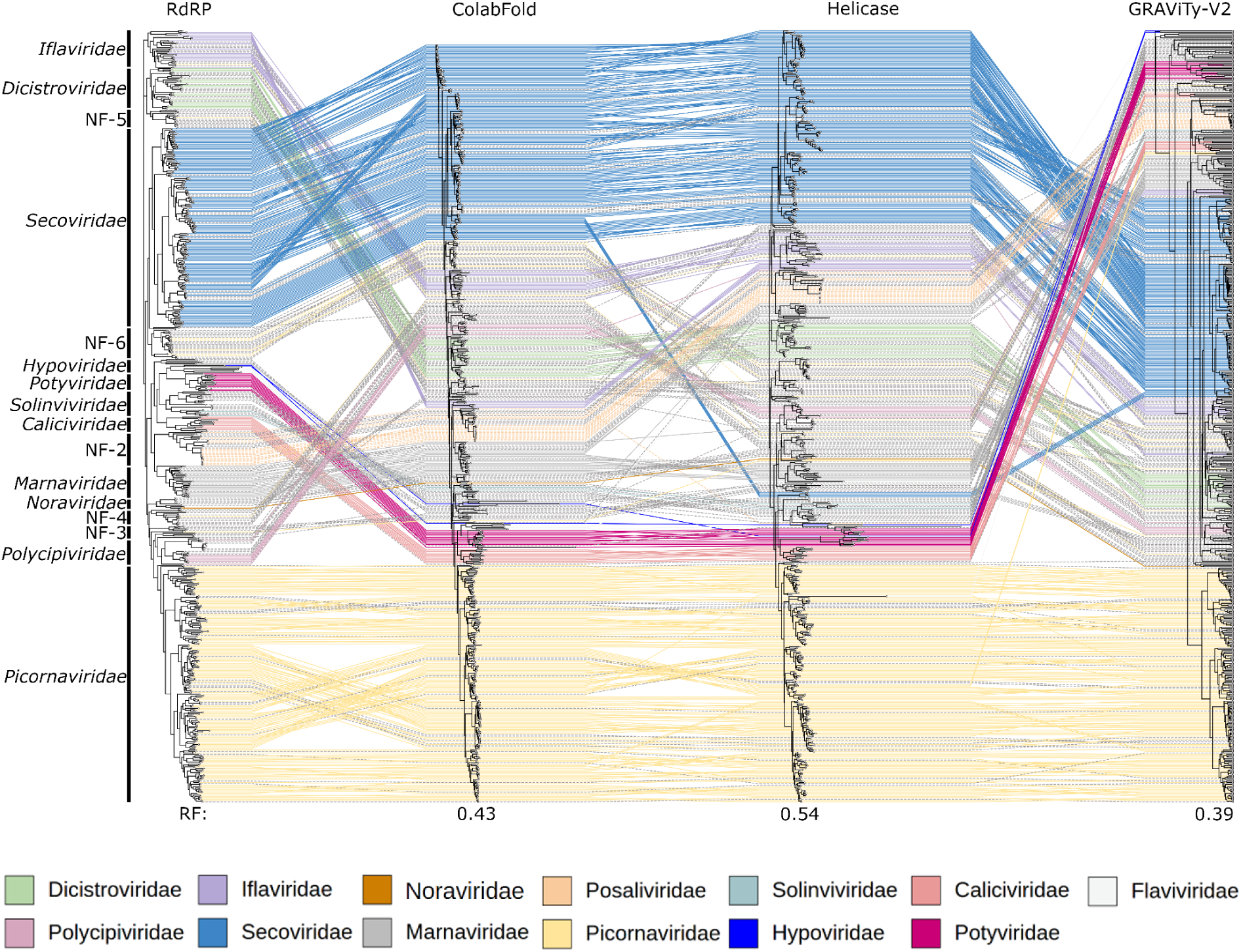
Comparison of phylogenetic trees highlights conservation of both existing and new family groupings across all analysis methods, but numerous genus-level rearrangements and, in the GRAViTyV2 phylogeny, inversions in relative depths of some families. Grey hashed lines indicate previously unclassified sequences. RF: Robinson-Foulds metric, calculated against the RdRp phylogeny.

**Table 2:**
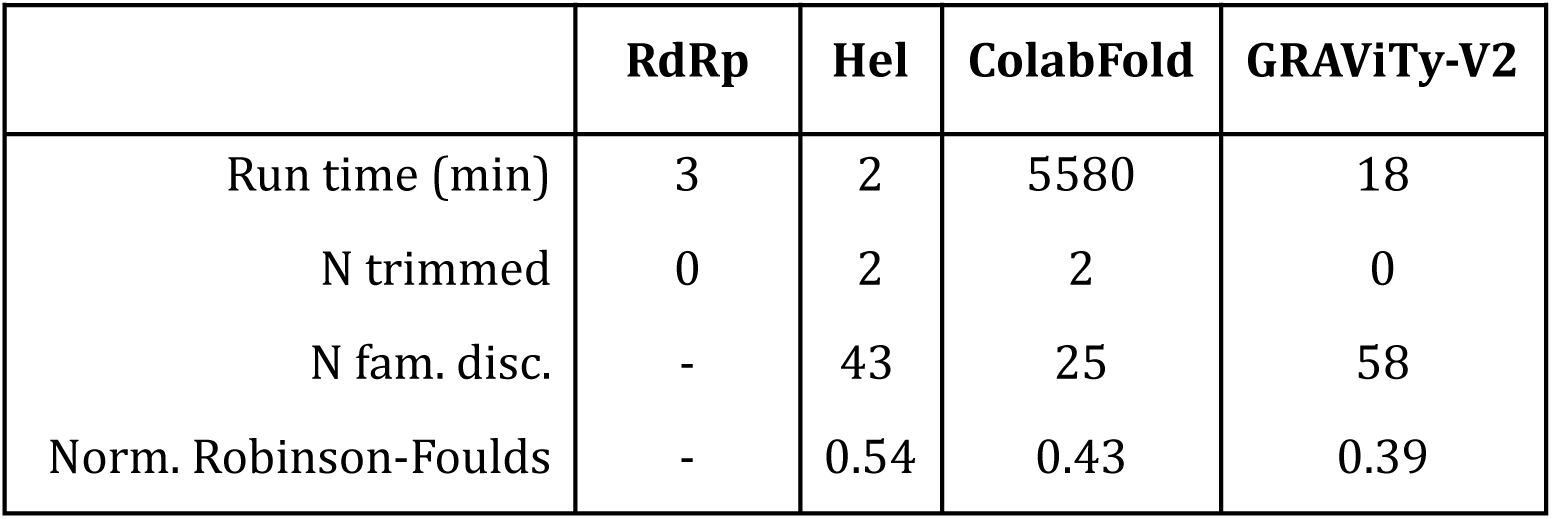
Summary statistics for each analysis method, comparing run time, number of sequences requiring trimming (N trimmed), number of family discrepancies versus RdRp sequence-based phylogeny (N fam. disc.) and Robinson-Foulds distance versus sequence-based RdRp tree.

|  | **RdRp** | **Hel** | **ColabFold** | **GRAViTy-V2** |
| --- | --- | --- | --- | --- |
| Run time (min) | 3 | 2 | 5580 | 18 |
| N trimmed | 0 | 2 | 2 | 0 |
| N fam. disc. | - | 43 | 25 | 58 |
| Norm. Robinson-Foulds | - | 0.54 | 0.43 | 0.39 |

Computing time for the complete pipeline was approximately 94 hours. Run times were comparatively short for both hallmark gene alignment and tree generation steps, at three minutes each. The GRAViTy-V2 analysis took approximately six times longer, and the structural prediction pipeline duration was nearly 2000 times longer. The long run times and comparatively high computing requirements (i.e. a discrete CUDA-enabled graphics card) for deep learning models such as ColabFold represent a key obstacle to their wider adoption and therefore necessitate a strong justification for their use in these contexts.

Phylogenetic placements for *Caliciviridae, Marnaviridae, Picornaviridae, Secoviridae*, as well as prospective family “Posaliviridae”, were generally stable between analysis methods, with few or no misclassifications or discrepancies (SI 2–3). Non-picorna-like virus families used as outgroup controls (*Hypoviridae*, *Potyviridae*, *Flaviviridae*) clustered appropriately in all methods except for in the Hel and GRAViTy-V2 phylogenies, where *Potyviridae* sequences were observed to partially cluster within *Picornavirales*, though inclusion of other ribovirian families in the second GRAViTy-V2 analysis caused them to group appropriately (section 3.5). Previously proposed subfamilies and genera within *Picornaviridae* [38] were well supported in all analyses, as were the existing genera within *Secoviridae*.

Classifications within *Solinviviridae, Polycipiviridae, Iflaviridae,* and *Dicistroviridae* were less stable between analysis methods. These families tend to exhibit greater variation in genome organisation, number of cistrons and host ranges in comparison with other families within *Picornavirales* [16, 21, 29, 38].

### 3.2 RdRp sequence-based analysis supports existing classifications and indicates new taxa

All previously classified genomes were placed within their appropriate family clusters in the sequence-based RdRp phylogeny, and each family-level grouping was supported by bootstrapping (≥ 0.7). Bootstrap support was similarly high for the majority of existing genus groupings. These observations highlight the value of RdRp phylogeny in ribovirian taxonomy, as well as the utility of automated RdRp domain extraction that was used in the UniViT pipeline. In addition to existing families in the *Picornavirales*, the analysis also revealed seven additional clusters whose branch depths were indicative of family-level groups (provisionally named ‘NF-1–7’, section 3.6), all of which were supported by bootstrap values of 0.76 or higher.

### 3.3 Hel phylogeny supports most classifications inferred from RdRp sequence-based analyses

The Hel phylogeny was concordant with the RdRp analysis regarding family placements for *Caliciviridae*, *Noraviridae*, NF-4 (*Macrobrachium rosenbergii* virus 10–13), *Picornaviridae* and *Iflaviridae*, and was highly similar (barring one or two outliers) for *Secoviridae, Dicistroviridae, Marnaviridae*, and NF-2 (“Posaliviridae”). Major differences were, however, observed for *Solinviviridae*, in which a presumptive new genus (*Solinvi-1*, *Riportus pedestris* virus 1) formed a new clade distinct from any family. Furthermore, a presumptive new family, NF-1, apparent in the RdRp phylogeny, was not reproduced in the Hel analysis, with members clustering with *Marnaviridae*. Similarly, both a presumptive new genus within *Polycipiviridae* (Polycipi-1) and the entire NF-3 clade either outgrouped or clustered with existing families on analysis of the helicase tree. Helicase domains within picornaviruses are considered highly conserved [35] and are known to make coherent phylogenies [11] but are however less conserved than RdRp domains and are additionally shorter. Picornavirus Hel domains are additionally prone to recombination, especially in pathogenic enteroviruses [33], which may further limit its usefulness in taxonomic investigations. These factors can make Hel phylogenies more challenging to interpret, and for this reason less reliable than counterpart RdRp sequence-based analyses.

### 3.4 Structural predictions follow RdRp

RdRp structural predictions for eight RdRP domain sequences failed (listed as “conversion failed” in SI 2) due to low pLDDT scores and were omitted from the analysis. Visual sense checks of molecular structure alignments confirmed that automated domain extraction produced biologically feasible structures, and that this was repeatable across the dataset (Fig. 4).

**Figure 4:**
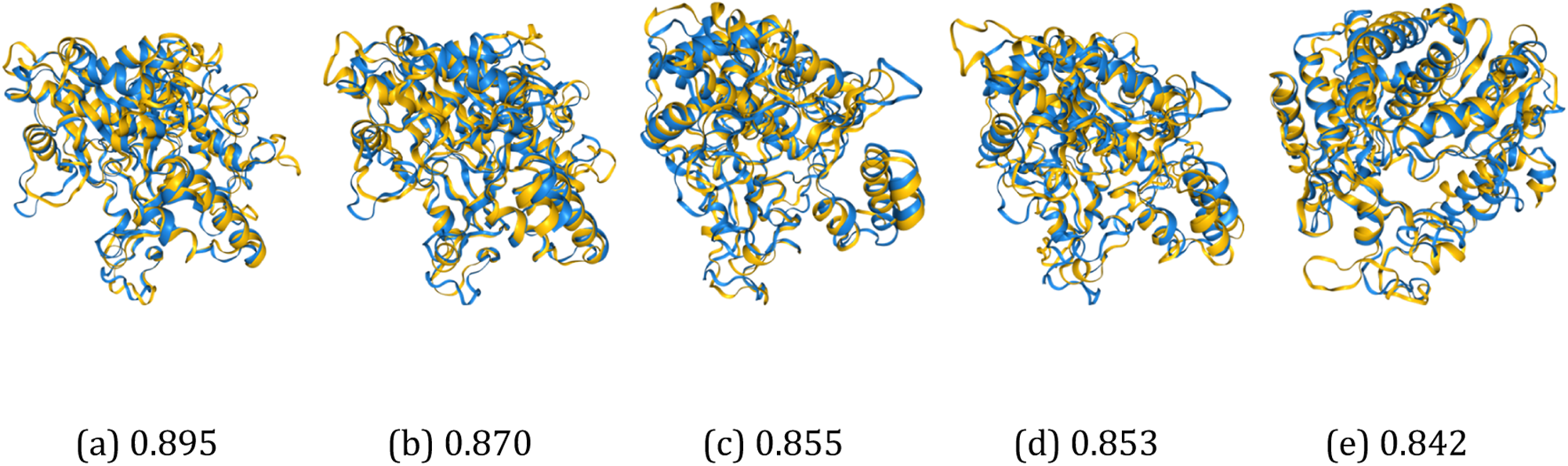
RdRp domain structural prediction comparisons between infectious flacherie virus (*Iflavirus flacherie*, blue) and random selections of structures across the dataset reveal biologically feasible tertiary structures with anticipated degrees of homology (by pLDDT score). (a) Deformed wing virus (*Iflavirus aladeformis*), RMSD: 3.14; (b) Macrobrachium rosenbergii virus 12 (NF-4), RMSD: 2.95; (c) Mobovirus A (*Picornaviridae*), RMSD: 3.27; (d) Mayfield virus 1 (*Noraviridae*), RMSD: 3.43; (e) Biomphalaria virus 1 (*Nora*-2), RMSD: 3.26.

Classifications for almost all family and genus groupings in the structural phylogeny were congruent with the RdRp sequence-based analysis, with noteworthy differences arising in only NF-1, *Polycipiviridae* and *Dicistroviridae*. As with the Hel phylogeny, the structure-based tree did not support the creation of NF-1, instead grouping all three viruses within *Solinviviridae*. The problematic sequences from *Polycipiviridae* and *Dicistroviridae* all clustered with other families (*Solinviviridae* and NF-5, respectively), in contravention of the RdRp phylogeny but consistent with Hel and GRAViTy-V2 analyses.

RdRp sequence-based phylogenies are generally expected to follow RdRp structural phylogenies, as protein structural conservation should be strongly correlated with sequence similarity. This correlation is however weakened in more distantly related protein structures [1]. Accordingly, the disagreements that were observed with sequence-based phylogeny arose in viruses that were also hard to place by other methods, indicating comparatively greater diversity in these clusters.

### 3.5 GRAViTy-V2 heatmaps reveal extent of picornavirus diversity

The GRAViTy-V2 phylogeny was the most divergent from the RdRp sequence-based phylogeny, though families *Marnaviridae, Picornaviridae, Secoviridae*, NF-2, NF-4, and NF-5 were largely concordant between the two approaches. In the majority of instances where there were violations, largely in *Solinviviridae, Polycipiviridae,* and *Iflaviridae*, violations tended to present as creating outgroup clusters, rather than placing discrepant genomes within other families. This behaviour is typical of how changes in genome composition and/or organisation, which are not reflected in conventional hallmark gene analyses, can influence composite Jaccard distances calculated by GRAViTy-V2, and highlights how an analysis method that is based on complete genome sequences can generate additional information over hallmark gene alignments.

Viruses within *Solinviviridae*, for example, may be either mono- or dicistronic, likely resulting from a ribosomal frameshift site between capsid and capsid extension proteins [2, 28, 31]. Accordingly, the dicistronic *Invictavirus solenopsis* cluster of solinviviruses formed an outgroup in the GRAViTyV2 analysis. Similarly, the GRAViTy-V2 analysis split *Polycipiviridae* into two clusters, as polycipiviruses within the *Sopolycivirus* cluster possess an additional, sixth ORF [13, 21]. While measuring cistronicity of a genome cannot in isolation be used as a method of producing taxonomic groupings, these data contribute to the depiction of genome organisational relationships through shared normalised PPHMM ratio pairwise comparison matrices, and PPHMM location ‘barcode’ graphs (SI 5).

GRAViTy-V2 output also revealed differences in genome organization through creating outgroups. *Iflaviridae* and *Dicistroviridae* were split into two clusters each. *Iflaviridae* are monopartite and functionally monocistronic with a single polyprotein strategy, whereas *Dicistroviridae* are monopartite but dicistronic, using dual IRES-driven translation to separately express functional and structural proteins. Members of both families may assume one of two genome organisations: ‘Type I’, wherein the genome is arranged (from 5’→3’) as {structural block → replication block}, and ‘Type II’, an inverted arrangement of {replication block → structural block}. In Type II genomes, the orientation of genes within the structural block may additionally be inverted. These examples further emphasise how analysis of complete genomes as part of a taxonomy workflow can uncover both structural and functional characteristics, which may be used to resolve discrepancies between other complementary methods and hallmark gene phylogenies in instances where no consensus is found.

When compared with other ribovirian families using GRAViTy-V2, the order *Picornavirales* was revealed to be monophyletic and showed a Jaccard score demarcation boundary that was comparable to those of other orders in the *Riboviria* (*>* 0.95; Fig. 5). Similarly, families within the *Picornavirales* were demarcated at comparable levels to those of families in other orders of RNA viruses (Jaccard distances of 0.70 — 0.95). The Jaccard distance metric therefore supported the classification of both new and existing clusters as families within *Picornavirales*, and differentiated them from groupings at lower or higher taxonomic ranks.

**Figure 5:**
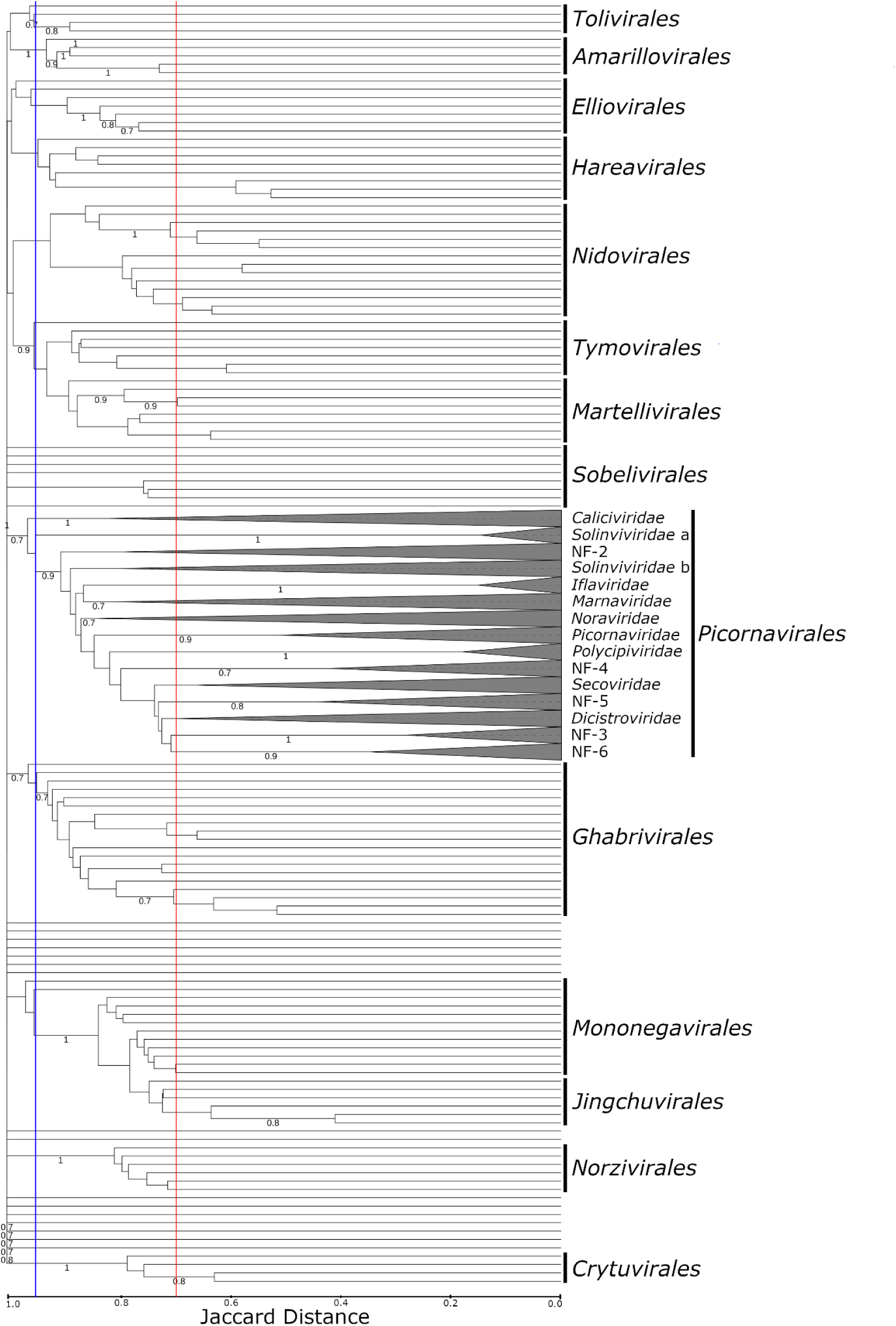
GRAViTy-V2 analysis reveals relative depth of both new and proposed *Picornavirales* clusters (*n* = 525) is consistent with the description of families within other representative ribovirian families (*n* = 135). Order and family lower demarcation boundaries are indicated at *>* 0.95 (blue) and *>* 0.70 (red), respectively All sub-family branches collapsed and outgroups (described in section 3.5) trimmed.

### 3.6 Description of suggested new taxa

A consistent finding between all analysis methods was the existence of six statistically supported clusters, labelled as presumptive new families (Table 3, SI 4). Divergence between these clusters was within a range of other families within other ribovirian orders (Fig. 5). Genome maps for exemplar members of each family (Fig. 6) revealed extensive diversity in genome length, content, and organisation between each grouping, consistent with their assignments to separate families. An expanded version of Table 1 to include new families is included in SI 6.

**NF-2** The corresponds to a previously proposed new family of posaviruses isolated from porcine and various other mammalian faeces, “Posaliviridae”. All members of the clade possessed monocistronic genomes and their grouping was strongly supported by all analysis methods.
**NF-3** 12 dicistronic or polycistronic metagenomically derived sequences from various plant and insect hosts. NF-3 was not supported by the Hel analysis, where all members either formed distinct outgroups, or clustered with *Noraviridae*.
**NF-4** A new grouping of four monocistronic *Macrobranchium rosenbergii* viruses, strongly supported by all analysis methods.
**NF-5** A cluster closely related to *Dicistroviridae* containing metagenomically derived viruses from insect-, bird-, bat- and rat-derived samples. NF-5 formed an outgroup in the GRAViTy-V2 analysis. Constituent genomes were fairly short, monocistronic, and several possessed zinc finger proteins detected via InterPro domain classification.
**NF-6** Cluster closely related to *Secoviridae* but derived from non-plant hosts. Four genomes (ON162339, PP818844, NC 033456, MG552116) that were placed within NF-6 in the RdRp phylogeny grouped with *Noraviridae* in the Hel phylogeny, and with NF-5 in the GRAViTy-V2 analysis. Genomes were monocistronic and possessed a complete {rhv-rhv-capsid protein} structural block.

**Figure 6:**
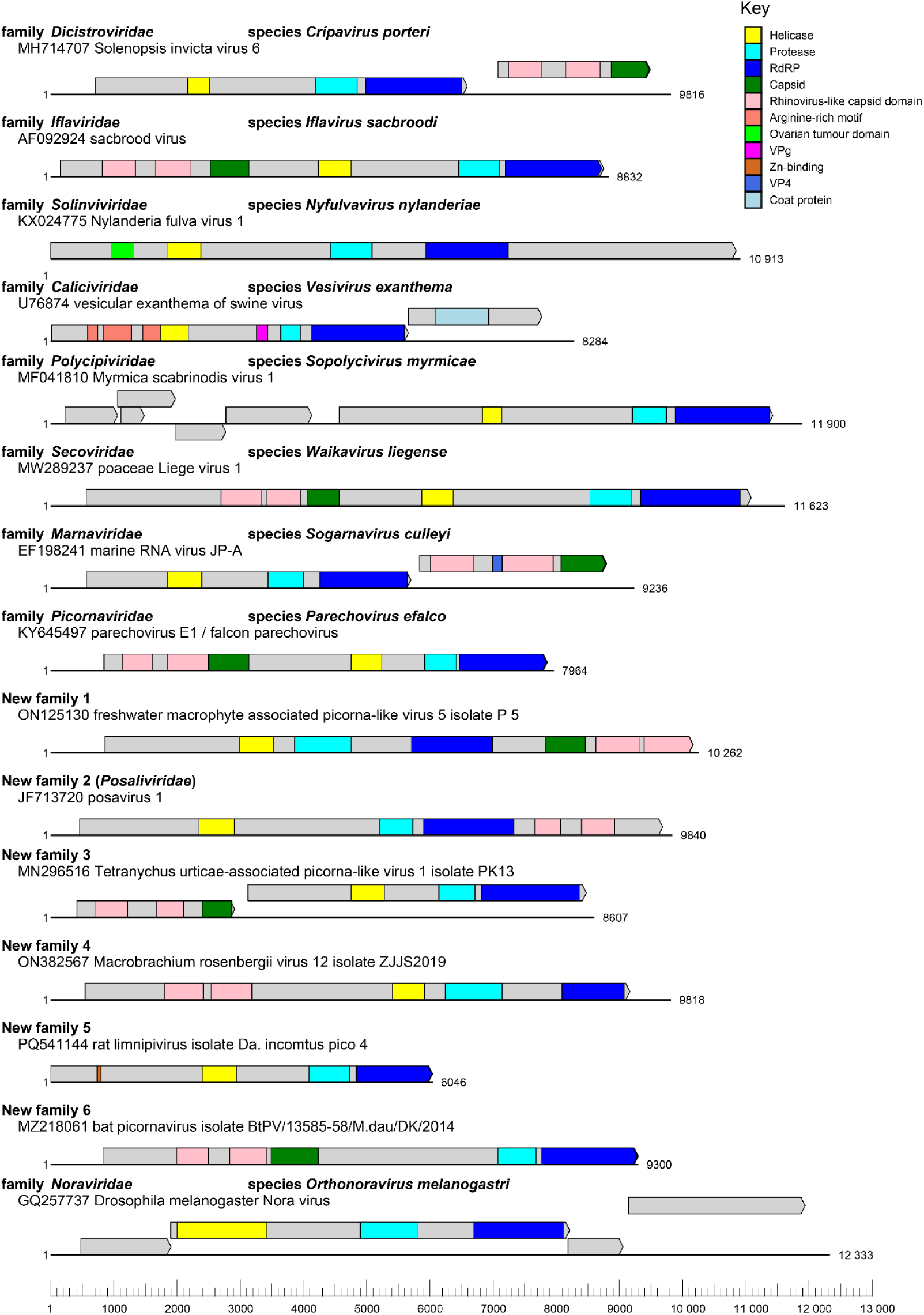
Representative single genomes from each classified and novel family within *Picornavirales*, drawn to scale, show extensive variation in the presence and organisation of protein-coding domains, as predicted by InterPro Scan. Additional protein coding domains may be present, but are too divergent from publicly available datasets to be detected with this tool.

**Table 3:** Summary of proposed new *Picornavirales* genera, grouped by family. *n* = number of genomes per genus.

| **Family** | **New Genus** | ***n*** | **Exemplar** |
| --- | --- | --- | --- |
| *Solinviviridae* | Solinvi-1 | 3 | KU958718 Riptortus pedestris virus-1 |
|  | Solinvi-2 | 10 | KP714071 Kilifi Virus |
| NF-1/*Solinviviridae* | NF-1-1<br>/Solinvi-3 | 3 | ON125130 Freshwater macrophyte associated picorna-like virus 5 isolate P 5 |
| NF-2 “*Posaliviridae*” | NF-2-1 | 1 | KR019688 Posavirus 3 |
|  | NF-2-2 | 1 | MN627463 Erysiphe necator associated picorna-like virus 2 |
|  | NF-2-3 | 1 | MH703051 Yongsan picorna-like virus 4 |
|  | NF-2-4 | 3 | MK606535 Posavirus 12 |
|  | NF-2-5 | 4 | LC131490 Picornavirales Bu-1 |
|  | NF-2-6 | 13 | JF713720 Posavirus 1 |
| *Noraviridae* | Nora-1 | 1 | MT416538 Bumble bee Mayfield virus 1-like |
|  | Nora-2 | 3 | KY024317 Biomphalaria virus 1 |
| NF-3 | NF-3-1 | 2 | MK533158 Tetranychus urticae-associated picorna-like virus 2 |
|  | NF-3-2 | 1 | MW314611 Linepithema humile picorna-like virus 4 |
|  | NF-3-3 | 9 | MG459012 Cherry virus Trakiya |
| NF-4 | NF-4-1 | 4 | ON382567 Macrobrachium rosenbergii virus 12 |
| *Polycipiviridae* | Polycipi-1 | 1 | KY024323 Biomphalaria virus 3 |
| *Picornaviridae* | Picorna-1 | 1 | MF370941 Mobovirus A |
|  | Picorna-2 | 1 | MN312220 Macaca fascicularis picornavirus |
| *Iflaviridae* | Ifla-1 | 2 | MW281483 Iflavirus achedomestici |
|  | Ifla-2 | 2 | AB000906 Iflavirus flacherie |
|  | Ifla-3 | 3 | AF323747 Iflavirus pernuda |
|  | Ifla-4 | 6 | AF092924 Iflavirus sacbroodi |
| NF-5 | NF-5-1 | 2 | ON382575 Macrobrachium rosenbergii virus 14 |
|  | NF-5-2 | 11 | MG662717 Blackbird arilivirus |
| NF-5/*Dicistroviridae* | Dicistro-1<br>/NF-5-3 | 2 | OQ599888 Asiatic toad picorna-like virus 1 |
| NF-6 | NF-6-1 | 3 | KY024322 Biomphalaria virus 2 |
|  | NF-6-2 | 17 | Bat picornavirus isolate<br>BtPV/13585-58/M.dau/DK/2014 |
| *Secoviridae* | Seco-1 | 2 | ON382570 Macrobrachium rosenbergii virus 15 |
|  | Seco-2 | 12 | AB009958 Satsuma dwarf virus |
|  | Seco-3 | 26 | OK040225 Cassava Torrado-like virus |
|  | Seco-4 | 3 | MH323435 Green Sichuan pepper nepovirus |
|  | Seco-5 | 6 | AY303786 Arabis mosaic virus |

An additional family-like grouping of three monocistronic viruses of freshwater hosts (NF-1; yellow catfish picornavirus (YCP) and two freshwater macrophyte associated picorna-like viruses (4, 5)), most closely related to *Marnaviridae,* was observed in the RdRp phylogeny. Although a single NF-1 member, YCP, did form a singleton family-like group in the structural tree, the other two sequences clustered with the solinviviruses in the ColabFold analysis. Conversely, the GRAViTy-V2 analysis placed YCP with three of the solinviviruses and the two freshwater macrophytes within a separate family-like grouping. The Hel phylogeny classified all three as marnaviruses. This highlights a rare instance of methodological disagreements with the gold standard method and represents an instance where expert supervision may be required to generate a definitive classification.

32 new bootstrap-supported sub-family level clusters were identified (Table 3), 29 of which were composed entirely of previously unclassified sequences. The remaining four clusters were identified by splitting *Iflaviridae*, which currently only contains a single genus, at branch depths comparable to genera in other families within the order. Presumptive new genera were proposed in instances where at least one virus formed a distinct group in at least two of the four analysis methods; where the cluster was split between methods, primacy was given to the RdRp phylogeny.

A potential genus apparent by RdRp phylogeny (SI 2), NF-5-3 in the family NF-5 contained two previously unclassified Asiatic toad picorna-like viruses. However, these sequences were grouped within *Dicistroviridae* (labelled alternatively as the new genus Dicistro-1) by the other analysis methods. As with family assignments, this further instance is another example where human input may be required to resolve discrepancies.

## 4 Discussion

The multimodal approach introduced in this study builds on our previous work in creating automated taxonomy workflows that are better suited to the current scale and pace of virus discovery. The majority of virus classifications in this analysis were comparable across the four methods used; in instances where there was disagreement, these corresponding sequences were automatically highlighted for further scrutiny. Genome organisational data from the full genomic analysis by GRAViTy-V2 was often able to reconcile discrepancies. While automated methods cannot replace manual classification by experts, we propose that the introduction of automated tools such as the UniViT pipeline may be used to perform an initial analysis of virus genome data, identify and align conserved gene blocks, and perform a range of comparison methods to guide taxonomic assignments.

While groupings were typically concordant between methods, there was a limited number of instances where all complementary methods disagreed with the RdRP hallmark gene phylogeny. This was observed once at the family level (NF-1), and several times at the genus level. In one instance, two sequences (Asiatic toad picorna-like viruses OQ599888, OQ599889) were placed in *Dicistroviridae* by RdRP sequence-based phylogeny, but within NF-5 in all other methods: these sequences were metagenomically assembled from lung and gut samples of Asiatic toads, and were originally labelled as ‘unclassified *Picornaviridae*’ [4] by maximum likelihood tree of RdRp sequences. Such samples are inherently challenging to assemble and classify, as it is frequently unclear whether the host organism was infected by, or had consumed a host for (dicistroviruses are insect-borne), the assembled virus. This further highlights how careful supervision by experts is required, and automated taxonomy workflows can help to realise greater efficiencies by expediting identification of problematic sequences.

While classical taxonomy workflows may not benefit substantially from incorporation of additional gene alignments or GRAViTy-V2 analysis, our results demonstrate that structural comparisons with ColabFold may be used to refine and support deeper clusters. Conversely, it should be recognised that a taxonomic classification is not a full evolutionary history of a virus. Beyond the RdRP hallmark gene, other genome components such as capsid genes may have entirely distinct evolutionary origins acquired through horizontal gene transfer (i.e. recombination and reassortment) that cannot be represented in the strictly hierarchical classification developed by the ICTV. Broader whole genome analysis, such as GRAViTy-V2, can be effective in identifying mosaicism genome modularity, and provide a potentially more complete account of a virus’ evolutionary history. Future work with UniViT will focus on the incorporation of further whole genome analyses methods and refining the pipeline to support its wider community adoption and its application for other RNA virus groups.

## Supporting information

SI 1: GRAViTy run files with dataset

SI 2: tabulated classifications

SI 3: summary of classifications

SI 4: rectangular trees (PDF)

SI 5: GRAViTy trees (PDF)

SI 6: Updated table of family descriptions

## 5 Acknowledgements

The authors extend their thanks and gratitude to Dr. Kentaro Tohma for their assistance in reviewing the technical content of this article.

## 6 Supplementary information

1. GRAViTy-V2 run parameters file and complete input data VMR.
2. Tabulated classifications.
3. Tabulated classification summary.
4. All rectangular trees in PDF format.
5. GRAViTy-V2 full output tables.
6. Expanded table 1 to show host range and genomic features of new families.

