## Supplementary figures and images for "Comprehensive hallmark gene sequence, genomic and structural analysis of *Picornavirales* viruses clarifies new and existing taxa"

### colabfold_rectangular.pdf

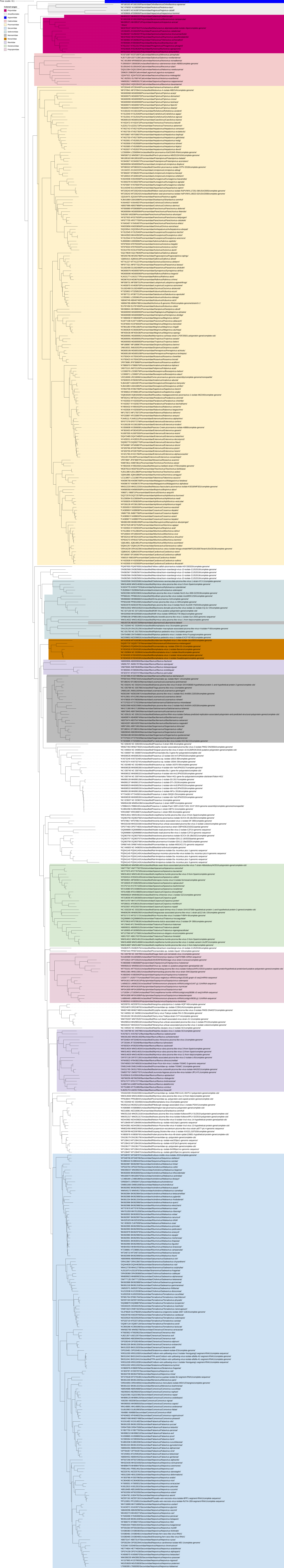

### GRAViTy_heatmap.pdf

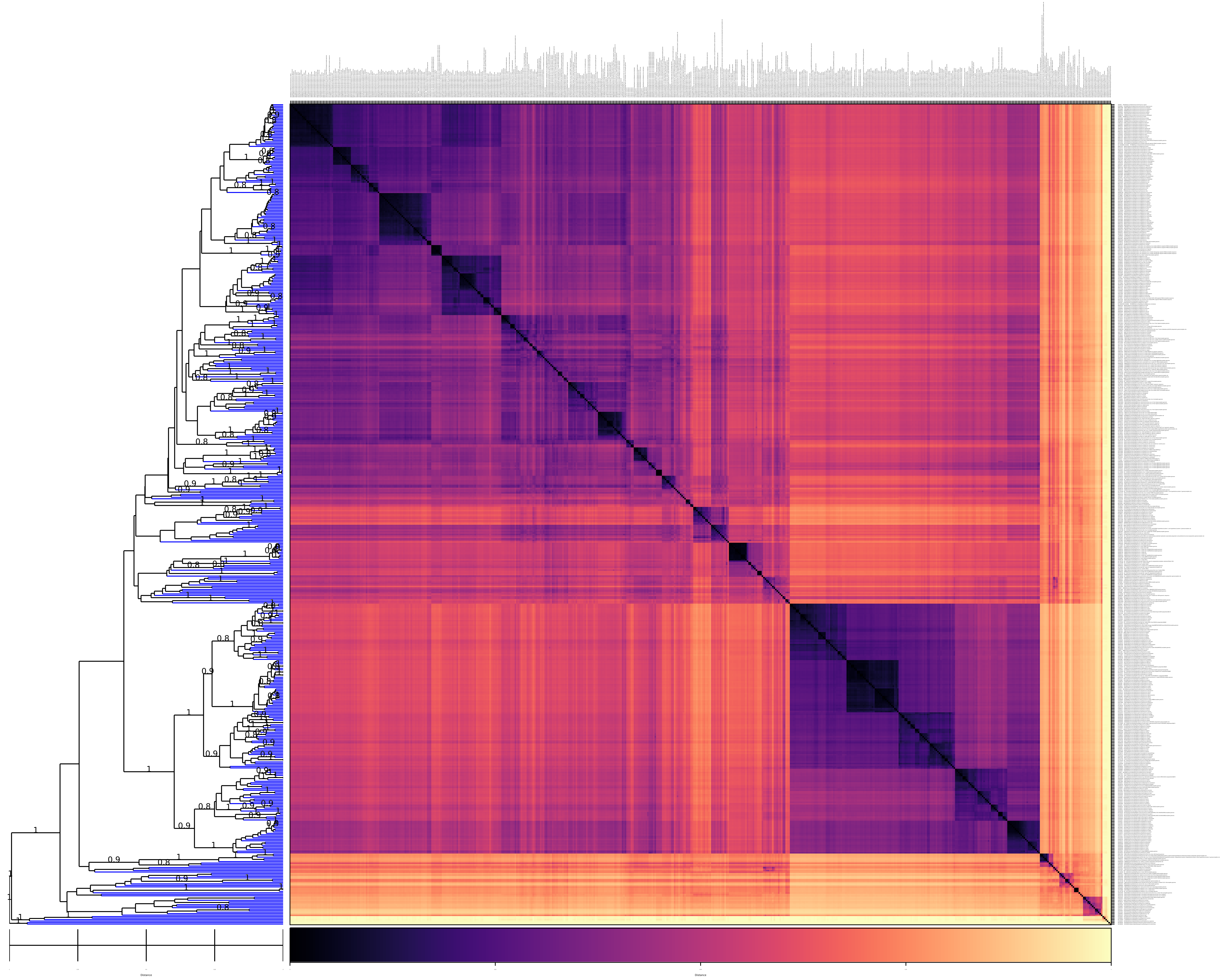

### gravity_rectangle.pdf

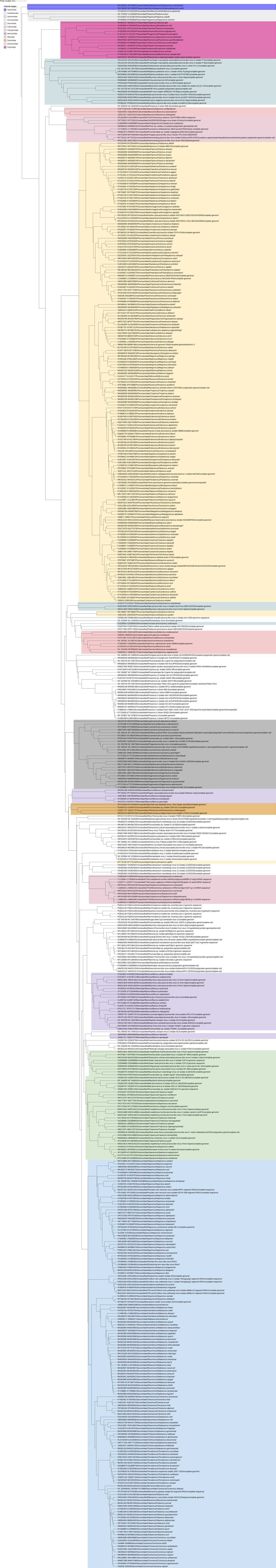

### helicase_rectangular.pdf

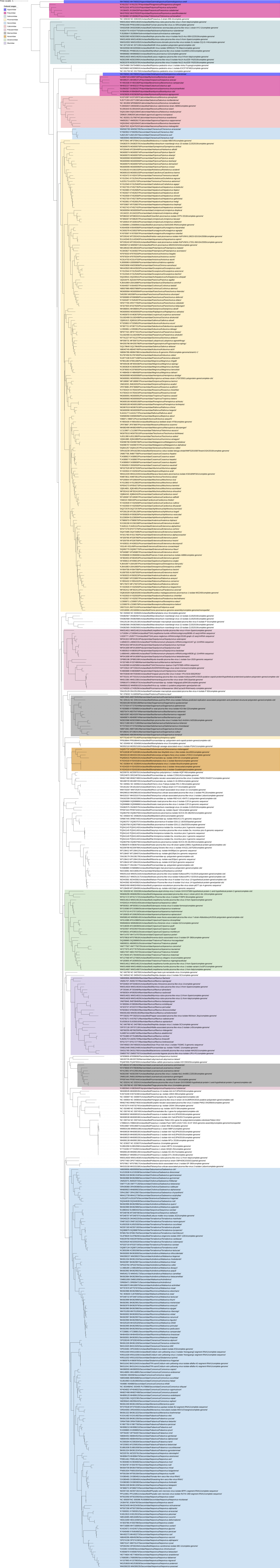

### Pphmm_locations.pdf

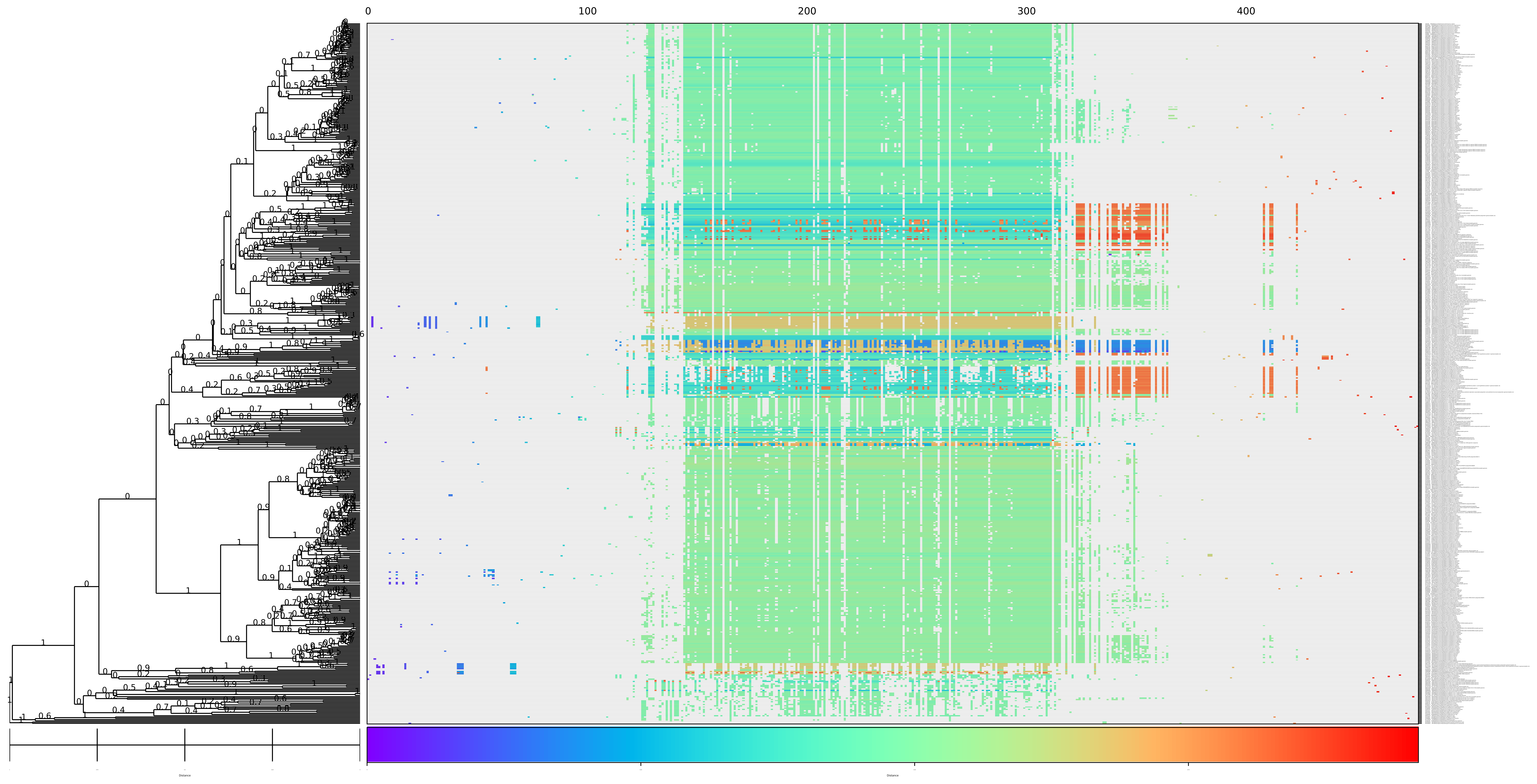

### rdrp_rectangular.pdf

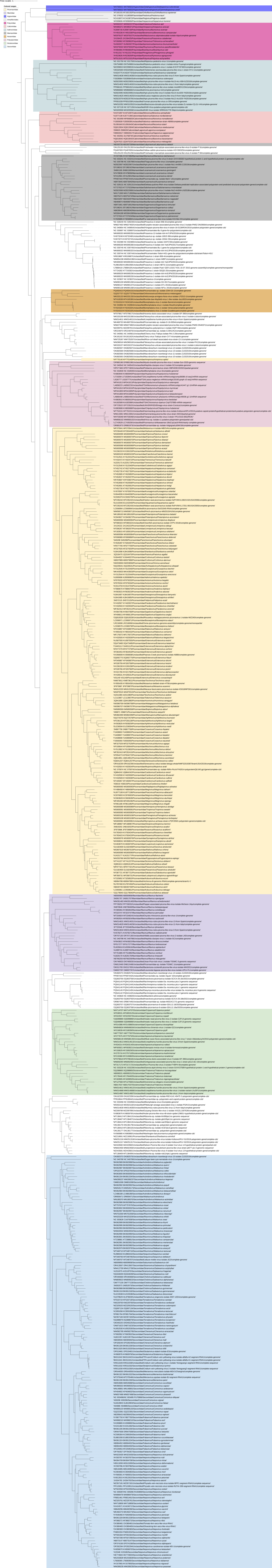

### Shared_norm_pphmm_ratio_matrix.pdf

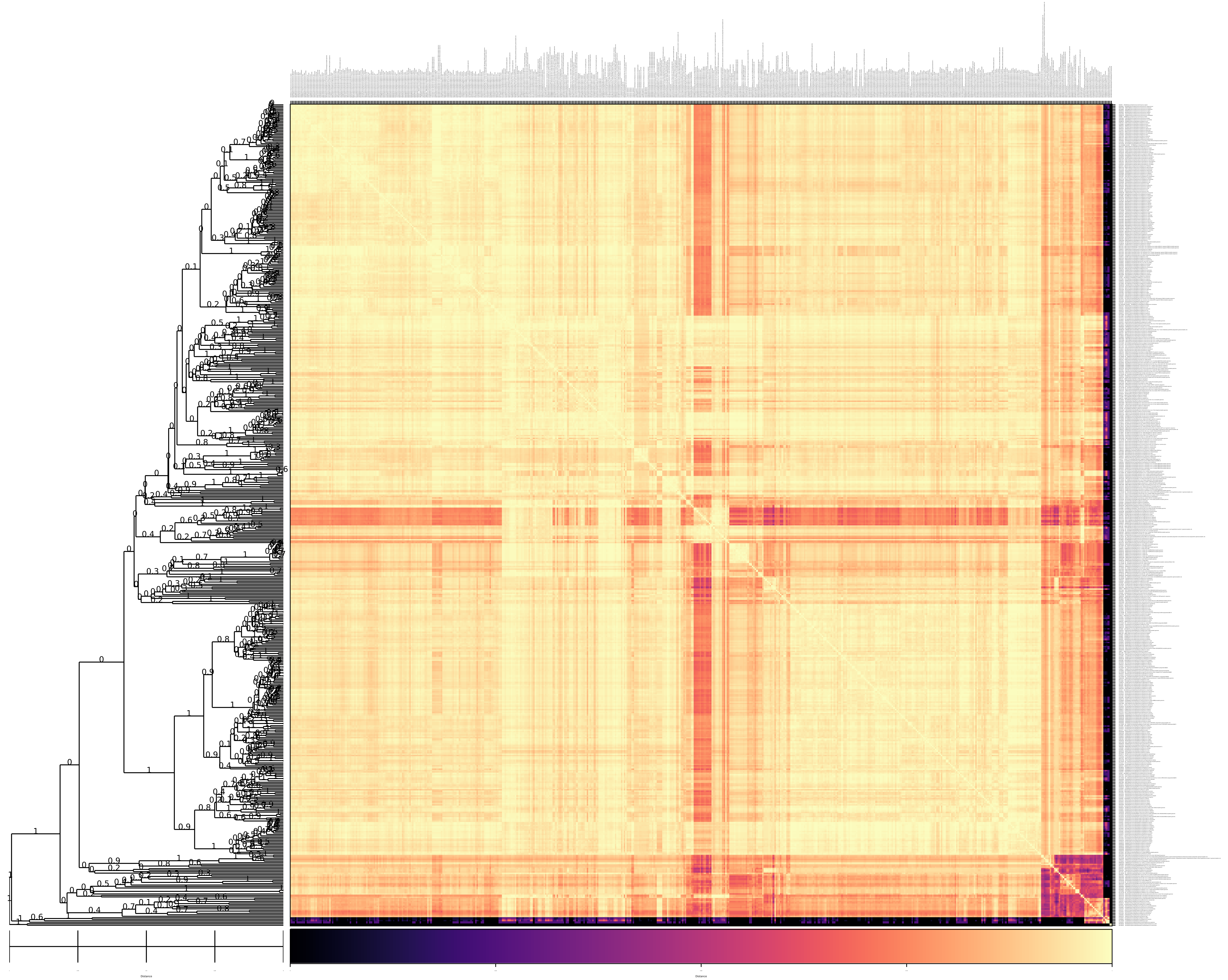
