## Supplementary material for "Comprehensive hallmark gene sequence, genomic and structural analysis of *Picornavirales* viruses clarifies new and existing taxa": SI 6: Updated table of family descriptions

Supplementary Information 6

| **Family** | **Host range** | **Genome** | **Ref** |
| --- | --- | --- | --- |
| *Caliciviridae* | *Mammalia* | 7–8 kb, monopartite polycistronic | [32] |
| *Dicistroviridae* | *Arthropoda* | 9–10 Kb, monopartite bicistronic | [29] |
| *Iflaviridae* | *Arthropoda* | 9–11 kb, monopartite monocistronic | [30] |
| *Marnaviridae* | Var. marine protists | 9–10 kb, monopartite monocistronic | [12] |
| *Picornaviridae* | *Animalia* | 7–10 kb, monopartite monocistronic | [35] |
| *Polycipiviridae* | *Arthropoda* | 10–12 kb, monopartite polycistronic | [21] |
| *Secoviridae* | *Plantae* | 9–14 kb, mono-/bipartite monocistronic | [6] |
| *Solinviviridae* | *Arthropoda* | 10–11 kb, monopartite polycistronic | [2] |
| *Noraviridae* | *Drosophila* | 12 kb, monopartite polycistronic | [7] |
| NF-1 | Freshwater macrophytes, yellow catfish (M) | 7-11 kb, monopartite monocistronic | - |
| NF-2 | *Animalia* | 10 kb, monopartite monocistronic | - |
| NF-3 | *Plantae, Arthropoda,* bat and parrot faeces (M) | 6-9 kb, monopartite bi-/polycistronic | - |
| NF-4 | *Arthropoda* | 10 kb, monopartite monocistronic | - |
| NF-5 | Bat, bird, rat and hedgehog faeces (M) | 8-11 kb, monopartite monocistronic | - |
| NF-6 | Sewage, platyhelminthes, molluscs, wasps, bat faeces (M) | 6-11 kb, monopartite polycistronic | - |

Table S6.1. Expansion of Table 1 in main article, to show range and genome organisation for newly proposed *Picornavirales* families. Host ranges for new families listed as sequence source, rather than confirmed infection. (M) = metagenomically derived
